# B cell receptor repertoire analysis unveils dynamic antibody response and severity markers in COVID-19 patients

**DOI:** 10.1101/2022.03.24.485649

**Authors:** Soumya Rao, Kriti Srivastava, Anupriya Verma, Achintya Das

## Abstract

Humoral and cell mediated immunity are critical against viral infections. The knowledge of composition, diversity, gene usage of the B cell repertoires helps in determining the immune response to SARS-CoV-2 infection. Examining B cell response provides insights on therapeutic antibodies, disease severity markers and aids in predicting vaccine response. We have analyzed public domain immunoglobulin sequencing data from PBMCs of SARS-CoV-2 infected individuals to gain a better understanding of B cell repertoire in patients. Public clonotypes showed increased usage of IGHV3, IGHV4, IGKV1, IGKV3, IGLV3 and IGLV2 family genes during the acute phase infection. Identical CDR3 sequences were identified for heavy (H), *kappa* (K) and *lambda* (L) chains across individuals, indicating the convergence of B cell selection during SARS-CoV-2 infection. While the immune repertoire dynamically changed over the course of convalescence, there were persistent clones across early and late timepoints. The diversity of antibody repertoire, measured by Shannon-Weiner diversity index for H and K chains, reduced during the acute phase of infection. In addition, the repertoire diversity was low in severe patients compared to patients with mild or moderate symptoms. Increased usage of *IGHV4-59* gene was observed in COVID-19 patients with severe symptoms requiring ventilator support at 2 weeks and 3 weeks post symptom onset. *IGHV4-59* is reported to have rheumatoid factor (RF) activity with high affinity for IgG and the elevated level of *IGHV4-59* provides a potential mechanism for the increased autoimmune responses in severe patients. Correlation of the clinical features with the B cell receptor repertoire dynamics elucidated public antibody clonotypes and disease severity markers for COVID-19.

## INTRODUCTION

Infectious diseases pose a major challenge to global public health. Infectious diseases caused by coronavirus family members have spiked three times in the population over the last 2 decades: severe acute respiratory syndrome coronavirus (SARS-CoV), middle east respiratory syndrome coronavirus (MERS-CoV) and recently, SARS-CoV-2. The acute pneumonia caused by SARS-CoV-2 infection was named “COVID-19” or “coronavirus disease 2019” by the World Health Organization (WHO). Since the emergence of SARS-CoV-2 in December 2019, the WHO has reported more than 462 million confirmed cases and 6 million deaths worldwide, with these statistics continuing to rise (World Health Organization, 18 March 2022). The rapid mutations leading to the emergence of newer variants such as B.1.1.529 (Omicron) have posed major concerns with increased infection rate and vaccine effectiveness (1).

While the initial symptoms of COVID-19 are mostly mild (fever, dry cough, fatigue), some patients gradually develop severe symptoms (e.g. dyspnea). In addition to causing lower respiratory tract pneumonia, SARS-CoV-2 also affects multiple organs such as the kidney, gastrointestinal tract, liver, brain and heart. While most people with mild or moderate COVID-19 symptoms recover after proper clinical care, the proportion of COVID-19 patients requiring hospitalization is high and they have an increased risk of admission to the intensive care unit (ICU) (2). This has caused a major burden on global public health resources. Some COVID-19 patients rapidly develop severe pneumonia, subsequently, multiorgan failure and death.

Humoral immune response plays a crucial defensive role in preventing SARS-CoV-2 infection. The virus SARS-CoV-2 targets cells in the nasal cavity and respiratory tract. The spike (S) protein of the virus is the principal antigen recognized by the protective antibody response against SARS-CoV-2. The S protein is cleaved into S1 and S2: S1 includes the receptor-binding domain (RBD) and the N-terminal domain (NTD); S2 contains the fusion peptide and heptad repeats (HR1, HR2) and mediates fusion with the host cell membrane (3). SARS-CoV-2 uses human angiotensin-converting enzyme 2 (ACE2) as an entry receptor through binding mediated by the RBD. The infection of the host cells is also mediated by transmembrane serine protease 2 (TMPRSS2). Pulmonary cells infected with SARS-CoV-2 release inflammatory signals that trigger the body’s antiviral immune response. SARS-CoV-2 infection may cause an excessive non-effective response and severe inflammation in some COVID-19 patients leading to “cytokine release syndrome” (CRS) (4). Despite availability of therapeutics for CRS, rapid progress of COVID-19 may cause some patients to miss the period when efficacious medication can be administered resulting in fatality.

In addition to CRS, it is also crucial to understand how adaptive immunity is established in COVID-19 patients in terms of B cell and T cell response. Knowledge of immune repertoire changes in COVID-19 patients would help in identifying disease severity predictive biomarkers, therapeutic monoclonal antibodies (mAbs) and in vaccine design. In older COVID-19 patients with comorbidities, diminished T cell repertoire and progressive defects in T cell and B cell function limited viral clearance and prolonged the innate proinflammatory response (4). An increasing number of mAbs have been isolated from convalescent COVID-19 patients and they have represented a promising intervention for COVID-19 disease (5). mAbs targeting RBD region of the S protein have received Emergency Use Authorization (EUA) in the United States.

BCR antibody clonotypes that share the same IGHV and IGLV genes among multiple individuals are called public antibody clonotypes. Public B cell clonotypes identified in the human antibody repertoires in response to diverse viruses, including Ebola, influenza, human immunodeficiency virus 1 (HIV-1), hepatitis C, respiratory syncytial virus revealed pattern of convergence of B cell clones with genetically similar antigen receptor genes in multiple individuals (6). Public clonotypes are of great interest in predicting the most common responses to vaccines in large populations as they provide understanding of viral epitopes that commonly induce antibodies in humans. The diversity of the antibody repertoire after virus infection depends on somatic hypermutation (SHM) and affinity maturation. Understanding molecular diversity and evolution of B cell receptor following SARS-CoV-2 infection helps elucidate the dynamics of the adaptive immune system within individuals. BCR-sequencing provides rapid and efficient methods for acquiring BCR heavy and light chain repertoires.

Analyzing complete repertoires provides a better understanding of the immune response to SARS-CoV-2. In this study, we analyzed the immunoglobulin repertoire of the COVID-19 patients to investigate the antibody response markers. We have identified novel public clonotypes, dynamic repertoire changes and disease severity markers during SARS-CoV-2 infection. Multiple studies have investigated the immunological features associated with SARS-CoV-2 infection. While most of the studies have focused on identifying neutralizing antibodies against the virus (7), some of them have also performed integrative analysis of BCR sequencing with the whole transcriptome (4). We employed comprehensive analysis of high-throughput BCR sequencing data from the peripheral blood mononuclear cells (PBMCs) obtained from COVID-19 patients in acute infection or convalescence (8). While the authors of the publication focused on identifying neutralizing antibodies against SARS-CoV-2, we have used the sequencing data to analyze the correlation of the antibody response with clinical features. We have assessed the antibody response with respect to public clonotypes, time-dependent changes and disease severity markers. The findings help in deciphering potential predictors for COVID-19 outcomes and novel potential mAb treatments for COVID-19.

## RESULTS

### Study design and analysis overview

The European Nucleotide Archive (<u>ENA</u>) was searched for bulk BCR sequencing datasets from COVID-19 affected individuals and 192 datasets (searched on 17 August 2021) were screened. SRA dataset PRJNA648677 was selected as 1) it was bulk BCR-seq experiment with data available to download and 2) data was available from chronological samples. The dataset possessed BCR sequencing data (separate libraries for heavy chain, *kappa* light chain and *lambda* light chain) derived from the PBMCs of individuals infected/recovered from COVID-19. While chronological samples were available for 7 patients (3 timepoints each for 2 patients and 2 timepoints each for 5 patients), 10 patients had samples derived from a single timepoint. 9 patients had mild symptoms (that did not require supplemental oxygen or ventilator), 6 patients had moderate disease that needed supplemental oxygen without requiring a ventilator and 2 patients required ventilator support (**Figure 1A**).

**Figure 1.**
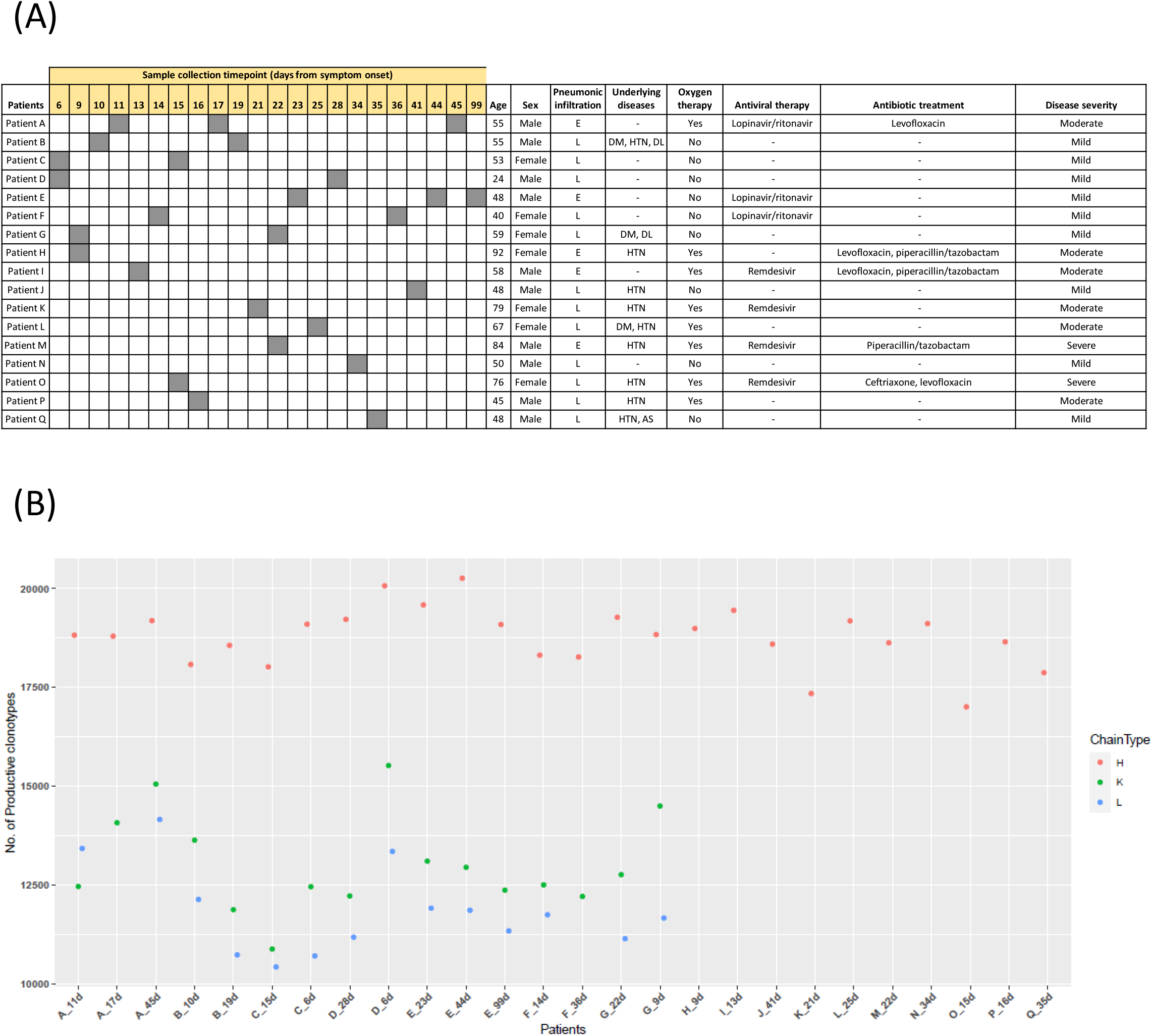
Study cohort and clonotyping. **A.** Demographic and clinical characteristics of study cohort. PBMCs were isolated from blood samples of the patients at indicated timepoints from symptom onset. DM, diabetes; HTN, hypertension; DL, dyslipidemia; AS, ankylosing spondylitis. Disease severity was defined as mild, moderate or severe based on the requirement of supplemental oxygen and ventilator. **B.** Productive clonotypes for H, K and L chains were identified from BCR-seq data for each patient. The raw sequencing data was preprocessed and clonotyping analysis was performed with annotations of V(D)J and CDR regions. While the samples A-to-G had data from heavy and light chain sequencing libraries, samples H-to-Q did not possess data have data on light chain K and L libraries.

While one (Patient D) of the 9 patients with mild symptoms had extensive pneumonic infiltration, 4 of the 8 moderate/severe patients had extensive pneumonic infiltration. The average age of mild patients was 47 years (24–59 years), with 50% of them having comorbidities such as diabetes (DM) or hypertension (HTN). 2 of the mild patients received antiviral therapy (lopinavir/ritonavir) and the others did not receive any antiviral or antibacterial therapy. The average age of moderate category patients (45–92 years) was 66 years and 4 of them had DM or HTN. The average age of patients in the severe category was 80 years, with both the patients having HTN. The gender distribution was similar between the mild and moderate/severe categories. While the severe patients received both antiviral and antibiotic therapy, there was varying degree of antiviral and antibacterial therapy among the moderate patients (**Figure 1A**).

Productive clonotypes for heavy chain (H), *kappa* light chain (K) and *lambda* light chain were identified for each of the patient samples (**Figure 1B**). Heavy chain clonotypes were higher with an average of 18,780 clonotypes compared to the light chain (K chain average = 13,032; L chain average = 11,837) (**Supplementary Table 1**). Light chain BCR-seq data was not available for patients H to Q.

### Public clonotypes during acute phase infection

Public clonotypes reveal the extent of convergence of B cell selection during an infection. The B cell selection is mediated by low-affinity recognition of viral surface antigens by germline-encoded naïve B cell receptors. Public clonotypes also aid understanding of viral epitopes that commonly induce antibodies in humans and thus have implications in predicting the most common responses to vaccines in large populations. Most efforts in characterizing public clonotypes in response to SARS-CoV-2 focused on neutralizing clones targeting RBD and NTD domains of S1 protein (2). However, public clonotypes directed to sites such as S2 domain are less characterized. Epitopes on the S2 domain may be of similar importance because of the increased sequence conservation across different strains of SARS-CoV-2.

To characterize the antibody response during the acute phase of infection, we analyzed the public clonotypes for H, K and L chains from samples with timepoints ranging from 9 days to 21 days. V segment usage analysis of clonotypes across the samples revealed higher frequency of IGHV3 family genes followed by IGHV4 family (**Supplementary Figure S1A**). At the gene level, there was increased usage of *IGHV3-23* (average 10.7% of the repertoire) and *IGHV3-30* (6.9%) genes across all the samples (**Figure 2A, top**). Among the IGHV4 family gene segments, *IGHV4-34* (5.5%) and *IGHV4-59* (5.2%) were predominant. CDR3 analysis for *IGHV3-23* showed the consensus amino acid sequence (**Figure 2A, bottom**). Stereotypic neutralizing antibody clonotypes were reported to be encoded by *IGHV3-53* and *IGHV3-66* V genes (8). These clonotypes were observed in our analysis, however with a lower frequency (1.45% and 0.8% respectively for *IGHV3-53* and *IGHV3-66*) during the acute phase infection. Among the light chains, there was increased frequency of IGKV1, IGKV3 family for kappa chain (**Figure 2B**) and IGLV3, IGLV2 for lambda chains (**Figure 2C**). For the kappa chain, increased frequency was found for *IGKV3-20, IGKV3-15, IGKV3-11, IGKV1-39* and *IGKV1-5* at the gene level (**Supplementary Figure S1B**). For L chain at the gene level, there is increased frequency of *IGLV3-19* followed by *IGLV2-14* (**Supplementary Figure S1C**).

**Figure 2.**
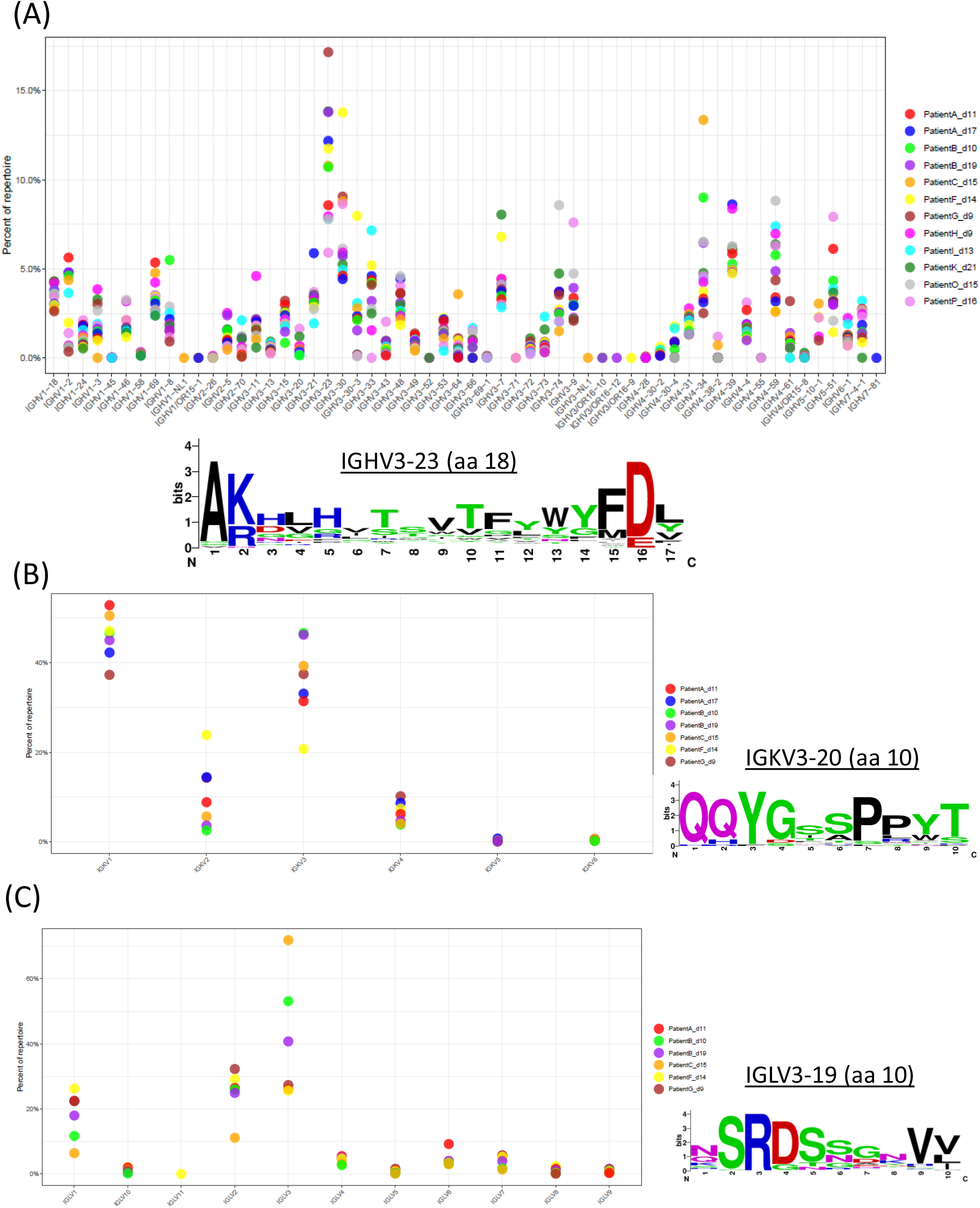
Gene and family usage of public clonotypes during acute phase infection. Productive clonotypes of the samples were used to calculate gene/family usage frequency. (**A**) IGHV gene usage (top) and multiple sequence alignment (MSA) of CDR3 motif (18 aa) of IGHV3-23 (bottom). (**B**) IGKV family usage (left) and MSA of CDR3 motif (10 aa) of IGKV3-20 (**C**) IGLV family usage (left) and MSA of CDR3 (10 aa) motif of IGLV3-19.

Further, we analyzed acute phase repertoire features at the level of CDR3. We identified clonotypes with identical CDR3 amino acid sequence across patients. For H chain we did not find any clonotypes with identical CDR3 across all the samples. Relaxing the cutoff to identify clonotypes across majority of the patients identified clonotypes present in at least 50% of the samples (**Supplementary Figure S2**). For K chain analysis, there were 8 samples (from 5 patients A, B, C, F, G) from acute phase infection. These samples possessed 876 common clonotypes with identical CDR3 sequences (**Figure 3A**). The majority of the CDR3s are 9-amino acid long and showed sequence conservation (**Figure 3B**). For L chain, data was present for 6 samples (from 5 patients A, B, C, F, G) and analysis showed 681 overlapping CDR3s (**Figure 3C**). The amino acid length varies with major proportion being 11, 10 and 9 amino acids. Top CDR3s showed moderate sequence variations (**Figure 3D**).

**Figure 3.**
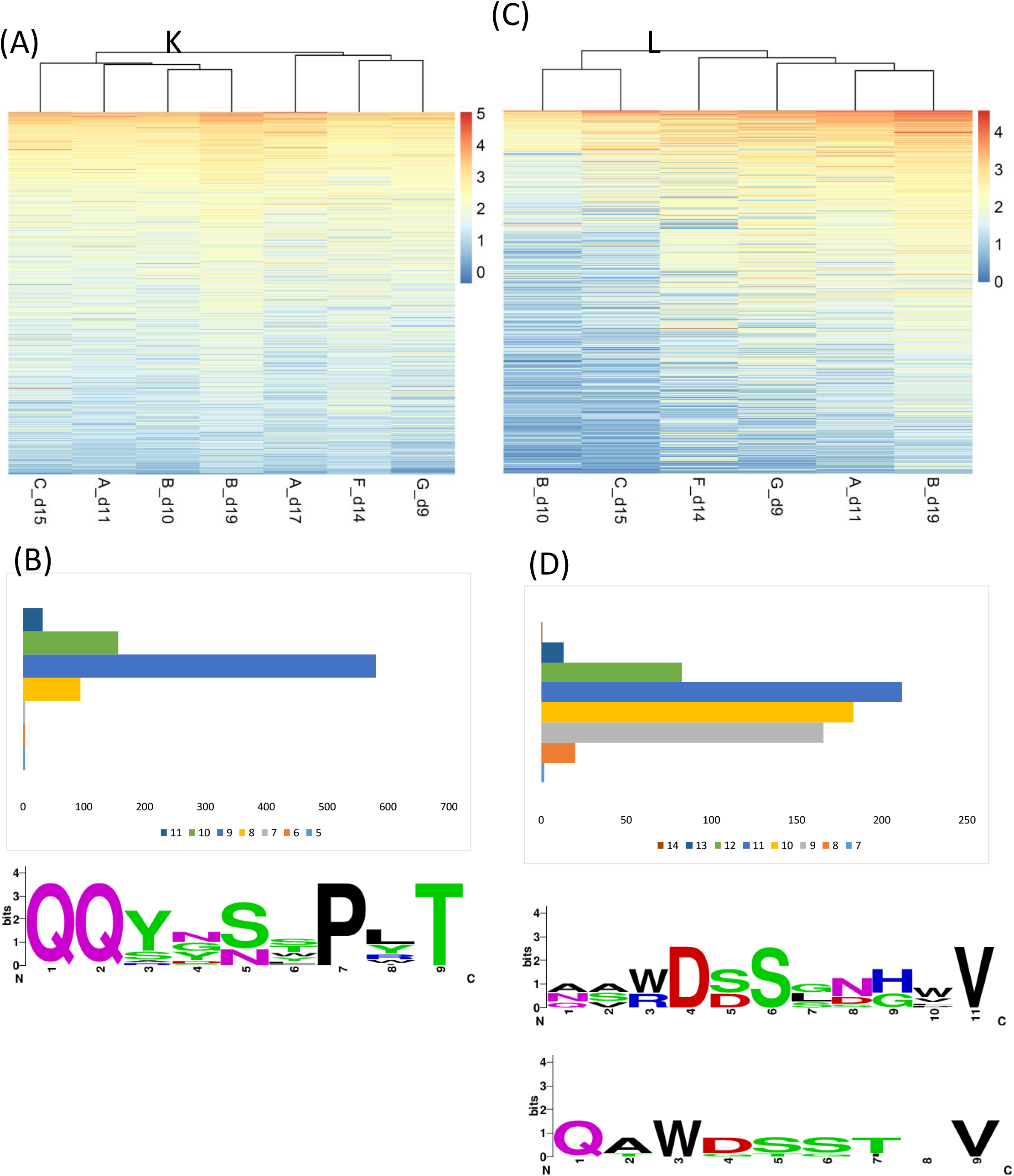
CDR3 sequence based analysis of public clonotypes during acute phase infection. Overlapping public clonotypes across samples were identified based on identical CDR3 amino acid sequence. The total number of sequences (normalized log2 read count) corresponding to the clonotype is represented as a heatmap for light chains K (**A**) and L (**C**). The amino acid sequence length of CDR3 for L and K public clonotypes, and the multiple sequence alignment of top sequences (median normalized read count >=2000) for K (**B**) and L (**C**) are represented.

### Dynamic antibody changes during course of SARS-CoV-2 infection and convalescence

Analysis of B cells with individual H, K and L chains can provide a complete view of antibody repertoire changes over time in the context of SARS-CoV-2 infection. We investigated how the immune response changes over the course of SARS-CoV-2 infection in each patient. For this analysis we considered patients A, B, C, D, E, F and G for whom chronological samples were available. Comparison of the repertoire at the CDR3 level between early and late timepoints for the same patient indicated that the overlap for H chain ranged from 4.5 – 9.2%, while for the light chains it ranged from 13.6 – 26.9 % (K) and 0 – 21.5% (L) (**Figure 4A**). The CDR3 overlap was the highest (H: 9.2%; K: 26.9%) for patient C between 6 days and 15 days from the symptom onset, while it was lowest for patient F (H: 4.5%; K: 13.6%) between 14 days and 36 days from the symptom onset (**Figure 4A**).

**Figure 4.**
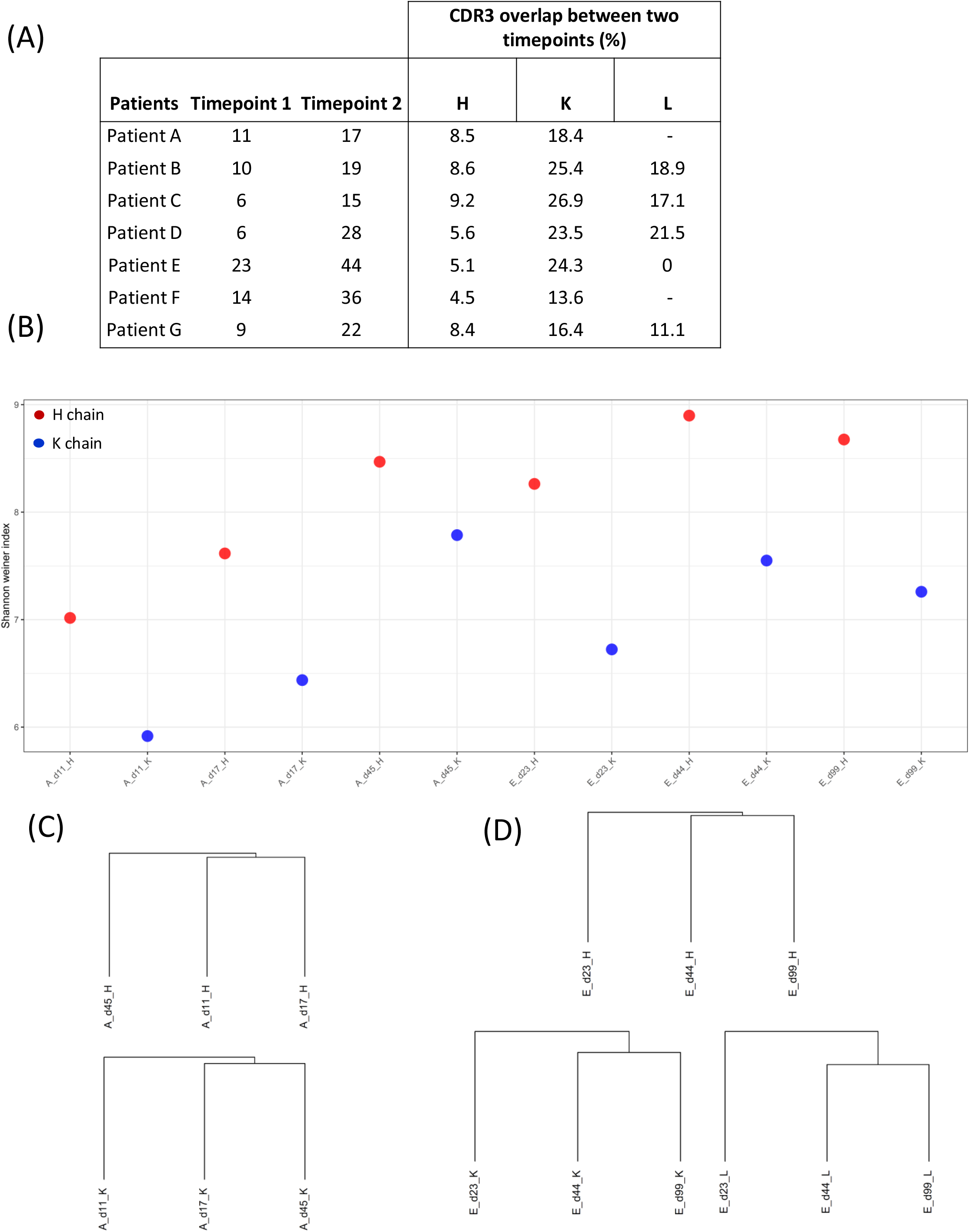
Dynamics antibody response: diversity. (**A**) The overlap in the antibody repertoire was investigated based on the presence of identical CDR3 sequence for each sample between the two timepoints. (**B**) Shannon Weiner index was calculated for Patient A and Patient E, the samples corresponding to 3 timepoints are represented. Jaccard similarity analysis of patient A (**C**) and patient E (**D**).

Further, we investigated samples A and E in detail. For both samples, data was available for 3 timepoints (patient A: 11, 17, 45 days; patient E: 23, 44, 99 days). The antibody diversity, as measured by Shannon-Weiner index, was low in the early timepoints for both H and K, increasing during convalescence (**Figure 4B**). Jaccard similarity analysis revealed that antibody repertoire of d11 and d17 were more similar compared to d45 for Patient A for both H and K chains (**Figure 4C**). For patient E, d44 and d99 were similar compared to d23 suggesting the influence of SARS-CoV-2 infection on the antibody repertoire in early stages of convalescence (**Figure 4D**).

#### Changes between week 2 to week 6

For patient A, 1.4% of the clonotypes overlapped between the timepoints 11, 17, 45 days for the H chain (**Figure 5A**). For K chain, the overlap was 7.7% (**Figure 5B**) (this comparison was not performed for L chain as clonotype data was not available for 17-day timepoint). Among the 1.4% overlapping H clonotypes (n = 726), read counts varied across the 3 timepoints. 2 distinct sets of clonotypes were observed. Set 1 clonotypes had a higher read count in the early timepoint (11 days) and this was sustained through 17 and 45 days (**Supplementary Figure S3A**). Read counts in Set 2 clonotypes were low in the early timepoints and became progressively higher at days 17 and 45. Among the overlapping K clonotypes (n = 2,444), proportion of Set 2 clonotypes were higher compared to Set 1 (**Supplementary Figure S3B**).

**Figure 5.**
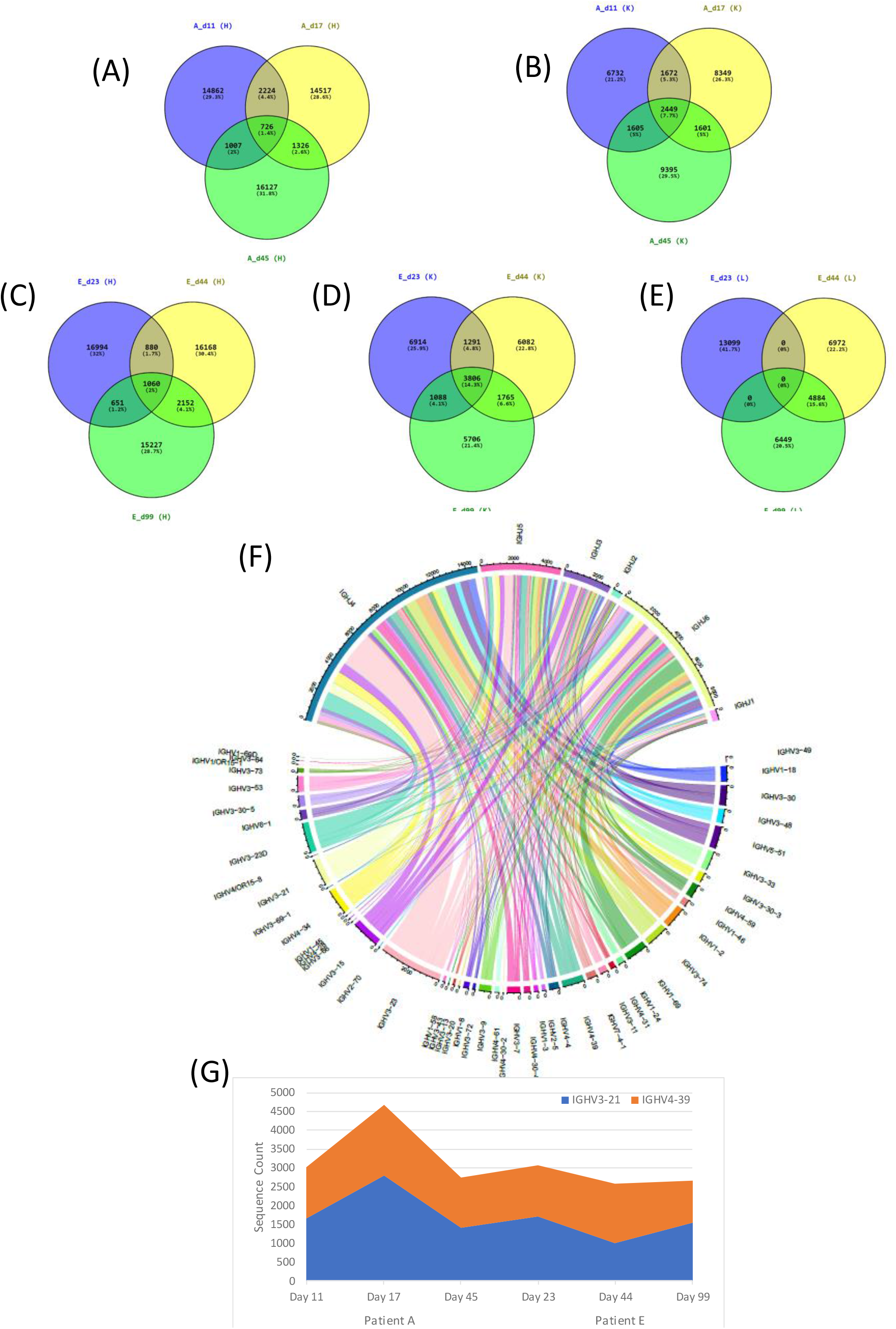
Dynamics of antibody response: overlapping clones. Venn diagram representing the overlapping clonotypes across timepoints for H chain (**A**) and K chain (**B**) for patient A and H (**C**), K (**D**) and L (**E**) chains for patient E. (**F**) Chord diagram representing V-J segment usage for patient A, day 11 sample. (**G**) Levels of IGHV3-21 clonotypes and IGHV4-39 clonotypes during infection and convalescence in patient A and patient E.

#### Changes between week 3 to week 14

For patient E, 2% H clonotypes overlapped between 23, 44 and 99 days (**Figure 5C**). The overlap was higher, 14.3% for the K chain (**Figure 5D**) and it was 0% for the L chain (**Figure 5E**). Heatmaps revealed a distinct pattern wherein majority of the high sequence count clonotypes at week 3 progressively had lower read counts at 44 and 99 days (**Supplementary Figures S3C and S3D**).

#### V-J segment usage analysis

V-J segment usage analysis revealed that segment usage of IGHJ4 is high compared to J5, J6, J3, J2 and J1 in all three timepoint samples for patient A (**Figure 5F and Supplementary Figures S3E and S3F**). *IGHV3-23* and *IGHV-23D* were predominant with similar frequencies across the 3 timepoints with most of them joining the J4 segment. *IGHV3-21* and *IGHV4-39* were present at day 11, peaked at day 17 and then reduced at day 45 (**Figure 5G**). For patient E, V-J segment usage is similar across 23, 44, 99 days (**Supplementary Figure S4A, S4B and S4C**). J4 is higher compared to J5, J6 and increases with time. While *IGHV3-23* was predominant, *IGHV3-21* and *IGHV4-39* levels were low (**Figure 5G**). For K chain, *IGKV1-39*, *IGKV1D-39*, *IGKV3-20* were predominant, and they remain at similar abundance across timepoints in both patients.

### Ig heavy chain gene-based disease severity marker

We compared the repertoire features of the patients with severe symptoms (patients M and O) with the patients with mild/moderate symptoms to identify antibody response markers that might correlate with disease severity. We chose two timepoints for analysis, 2 and 3 weeks.

We observed a specific pattern of increased V gene segment usage, *IGHV1-3*, *IGHV1-46*, *IGHV1-8*, *IGHV3-48*, *IGHV3-74*, *IGHV3-9*, *IGHV4-39*, *IGHV4-55*, *IGHV4-59* in severe patient O compared to the mild/moderate patients C, F and P at 2 weeks (**Figure 6A; Supplementary Figure S5**). The usage of *IGHV3-30* was low in severe patients compared to the mild/moderate individuals. The decreased usage of V3-30 is evident at 3 weeks in severe patient M compared to the mild/moderate patients E, G and L (**Figure 6B**). The repertoire diversity, measured by Shannon-Weiner index, is low for severe patient at 2 weeks compared to mild and moderate patients (**Figure 6C, top**). At 3 weeks, the repertoire index is low for both moderate and severe patient compared to the mild samples (**Figure 6C, bottom**).

**Figure 6.**
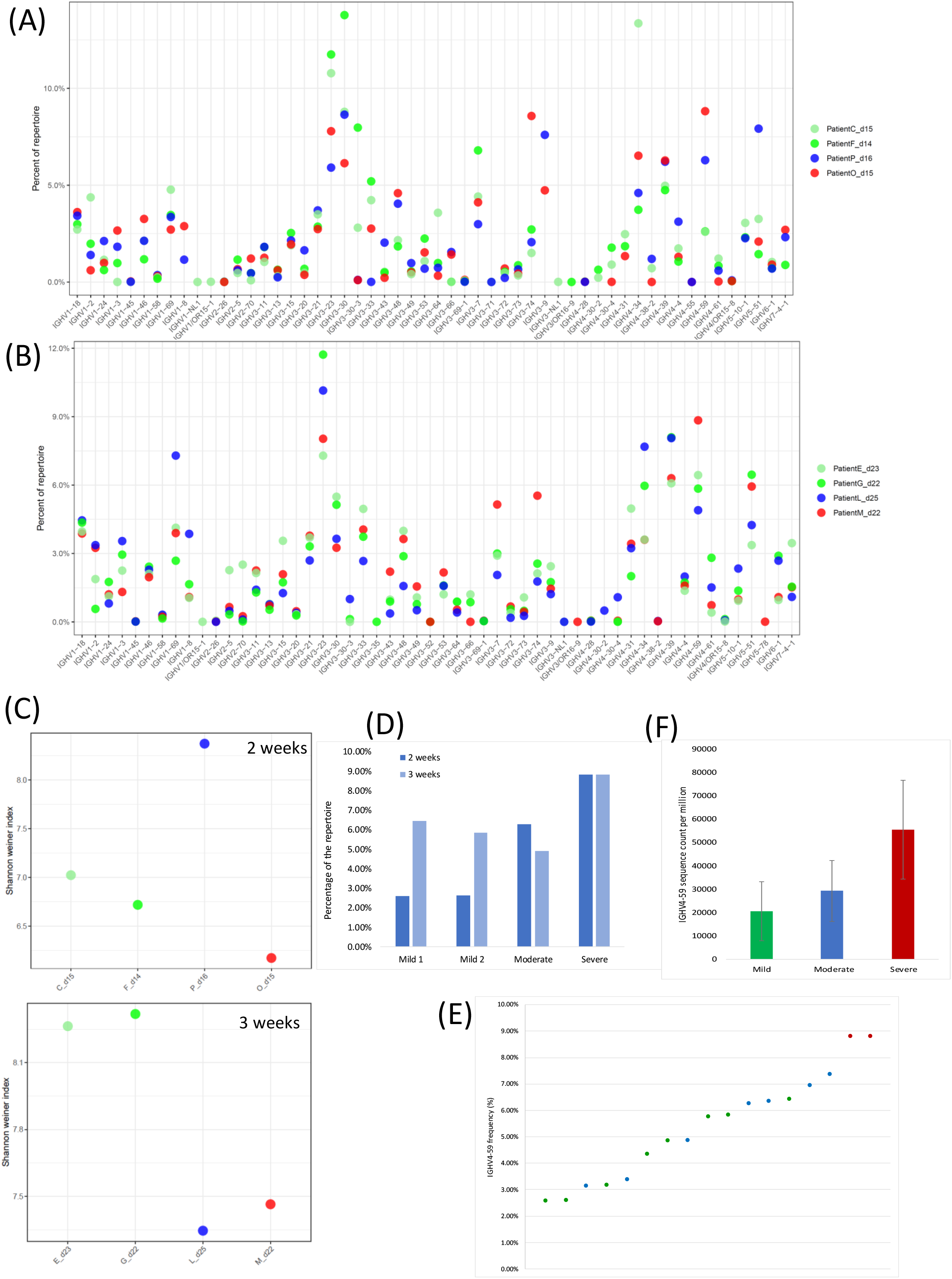
Antibody response markers in mild, moderate and severe patients. Repertoire V gene usage frequency at 2 weeks (**A**) and at 3 weeks (**B**) in mild, moderate and severe patients. Antibody repertoire diversity as measured by Shannon-Weiner index is represented at 2 weeks and 3 weeks (**C**), Gene usage of *IGHV4-59* (**D, E**) and sequence count is represented for mild, moderate and severe patients.

Closer examination of the V-J segment usage revealed increased usage of *IGHV4-59* in severe patients compared to mild/moderate patients at 2 weeks and 3 weeks (**Figure 6D**). The frequency of *IGHV4-59* across all patient samples reconfirmed the observation (**Figure 6E**). In addition, the average sequence count was lower in mild (20,363) and moderate (29,215) compared to the severe patients (55,480) (**Figure 6F**). IGHV4-59/IGHJ5-encoded antibody was expressed in several types of B cell lymphomas, and they were found to bind to IgG-Fc with high affinity. They were reported as rheumatoid factor (RF) (9), with IgG as major autoantigen, playing an important role in the pathogenesis of B cell lymphomas (10).

## DISCUSSION

We performed a systematic search to identify BCR sequencing datasets of COVID-19 patients and investigated key immunological features of humoral response against SARS-CoV-2 infection. We have performed comprehensive analysis of the data to decipher new insights including public clonotypes, repertoire dynamics and disease severity signature.

The analysis of the repertoire to identify public clonotypes during acute phase infection revealed enhanced usage of IGHV3 and IGHV4 family genes. *IGHV3-53* (1.45%) and *IGHV3-66* (0.8%) were observed at a lower frequency compared to *IGHV3-23* (10.7%) and *IGHV3-30* (6.9%) during acute phase infection. Increased usage of *IGHV3-23, IGHV3-30, IGHV4-34* and *IGHV4-59* were also reported in the public clonotypes for SARS-CoV-2 (6). Light chain clones with increased frequency of *IGKV3-20, IGKV3-15, IGKV3-11, IGKV1-39, IGKV1-5, IGLV3-19, IGLV2-14* were also concordant with the previous study.

Since naïve B cells undergo somatic hypermutations, clonal selection and class switching after antigen exposure, we examined the chronological events that occurred in all H, K and L clonotypes across patient samples. At the CDR3 amino acid level, 4.5-9.2% of the H chain repertoire overlapped between chronological samples while the overlap was 13.6-26.9% for K and 0-21.5% for L chains. The overall antibody diversity was lower at the early timepoint compared to the late timepoint for both patient A and patient E.

While we have uncovered B cell adaptations from our analysis, it is limited by fewer patient samples in the severe category and requires confirmatory studies from wider datasets. For severe patients, only heavy chain data was available. In addition, data from healthy control is not available and hence any comparison between the uninfected and infected/convalescent could not be performed.

Diversity was reduced in severe patients compared to mild/moderate at 2 and 3 weeks post symptom onset. Similar observations have been reported 4 weeks after post hospital discharge (11). Mor *et al* demonstrated that severe disease group had higher anti-SARS-CoV-2 receptor binding domain (RBD) plasma IgG titers (12). Correlation of antibody titer with the disease severity was not observed in the current cohort (8). Our analysis revealed the increased usage of *IGHV4-59* gene in patients with severe manifestations compared to those with mild/moderate symptoms. *IGHV4-59-encoded* antibodies are identified in lymphomas as rheumatoid factors with auto-immune activity. Enriched *IGHV4-59* in severe patients suggest potential increase in the autoimmune response in severe patients compared to the mild/moderate individuals. Mild binding activity was observed with IGHV4-59 encoding antibodies during convalescence against HCoV Spike protein, SARS2 S non-RBD, SARS2 RBD and ORF8 proteins (13). IGHV4-59/IGKV4-1 antibodies were observed during convalescence and reported to show binding activity against SARS-CoV-2 nucleocapsid protein (NP) (14).

The results of this study provide a comprehensive view of the B cell response in COVID-19 patients during acute phase infection and convalescence. We have characterized the public clonotypes in response to SARS-CoV-2 infection highlighting the preferential usage of subset of V and J genes. We have provided a detailed view of B cell receptor dynamics during infection and recovery phases. Our study also provides insights on immunoglobulin-gene based disease severity marker that can be of value as an early predictor of disease severity, helps patients with appropriate therapeutic intervention.

## METHODS

### Study design

This study was designed to investigate antibody response with SARS-CoV-2 infection from the BCR sequencing data. Systematic search of the BCR-seq datasets was undertaken and <u>PRJNA648677</u> was shortlisted based on predefined inclusion/exclusion criteria. The study cohort contained twenty-six blood samples that were collected from 17 SARS-CoV-2 positive patients. All 17 patients were confirmed to be infected by SARS-CoV-2 by a positive RT-qPCR result. Detailed clinical information was available (**Figure 1A**). PBMCs were isolated from blood samples and BCR sequencing was performed on an Illumina MiSeq platform as described before (8). While data was available for H, K and L chains for samples A-L, for the rest of the samples, only H library information is available.

### Clonotype annotation of NGS data

Paired end raw sequencing data was downloaded from SRA (<u>PRJNA648677</u>). FastQC was performed on read 1 (R1) and read 2 (R2) separately. The R1 and R2 were then assembled using pRESTO, The Repertoire Sequencing Toolkit (15). The reads were first assembled *de novo.* If *de novo* assembly failed, reference guided assembly was performed using V segment reference sequence from IMGT (the international ImMunoGeneTics information system). The assembled reads were subjected to trimming using quality Phred score 20 as cutoff. <u>Seqtk</u> (16) was used to convert sequences in FASTQ format to FASTA format. The quality trimmed reads were further processed using IgBLAST (17). IgBLAST enabled annotation of germline V, D and J genes, rearrangement junctions, delineation of framework and CDR boundaries. Human germline sequences for heavy, *kappa* and *lambda* from IMGT were used for VDJ and CDR annotation. The resultant file contained information such as the read count, frequency, CDR3 sequence, and V/D/J gene for each clonotype. Clonotype file was further processed to retain only productive rearrangements by filtering out the clonotypes with stop codon or out-of-frame rearrangement. Clonotypes with identical CDR3 were merged.

### Gene/family usage frequency plots

Gene and family usage quantification was performed by the R package *alakazam* (18). The relative abundance of V genes or families across samples was quantified by sequence frequency and visualized.

### Data visualization

Heatmaps were created using the R package <u>*pheatmap*</u> (19). R package *circlize* (20) was used to create chord diagram for visualizing V-J usage frequencies.

### Diversity

The Hill diversity index (qD) (21), which measures diversity in a population, was calculated for the set of BCR clones to determine overall repertoire diversity. R package <u>*vegan*</u> (22) was used diversity analyses. Shannon-Weaver diversity index (H’) was calculated on the clonotype file for each sample as follows:

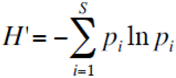

Where *p_i_* is the *relative abundance* of species *i, S* is the total number of species present, and ln is the natural logarithm.

The evenness was calculated as Pielou’s evenness J = H’/ log(S).

To assess the closeness of the repertoire across samples, Jaccard distance was calculated using vegan package. Jaccard distance is the number of species shared between two sites divided by the total number of species in both sites.

## Supplementary Figures

**Figure S1.**
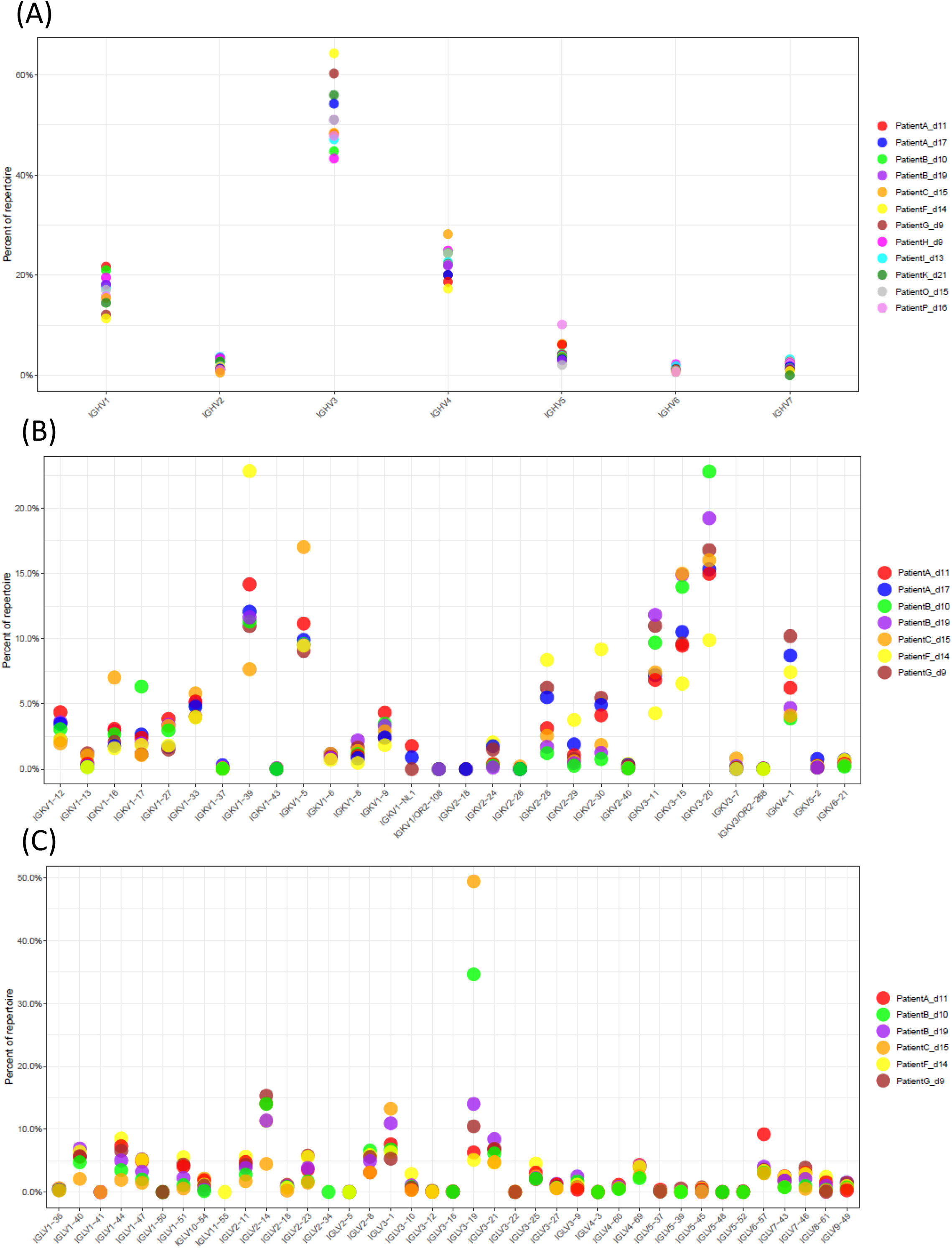
V gene/family usage of public clonotypes during acute phase infection. Productive clonotypes of the samples were used to calculate gene/family usage frequency for (**A**) IGHV, (**B**) IGKV and (**C**) IGLV

**Figure S2.**
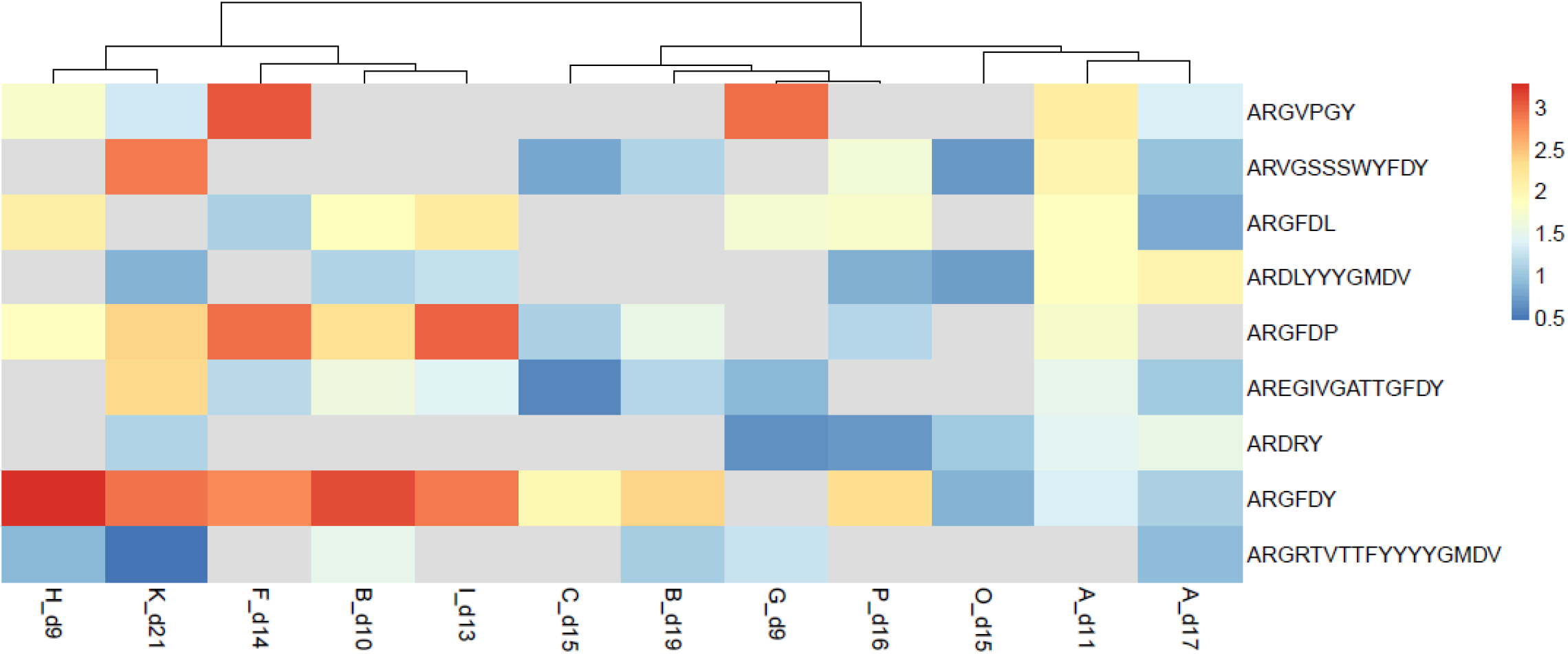
HCDR3 sequence based analysis of public clonotypes during acute phase infection. The overlapping public clonotypes across samples were identified based on identical CDR3 amino acid sequence. The total number of sequences (normalized log2 read count) corresponding to the clonotype is represented as a heatmap.

**Figure S3.**
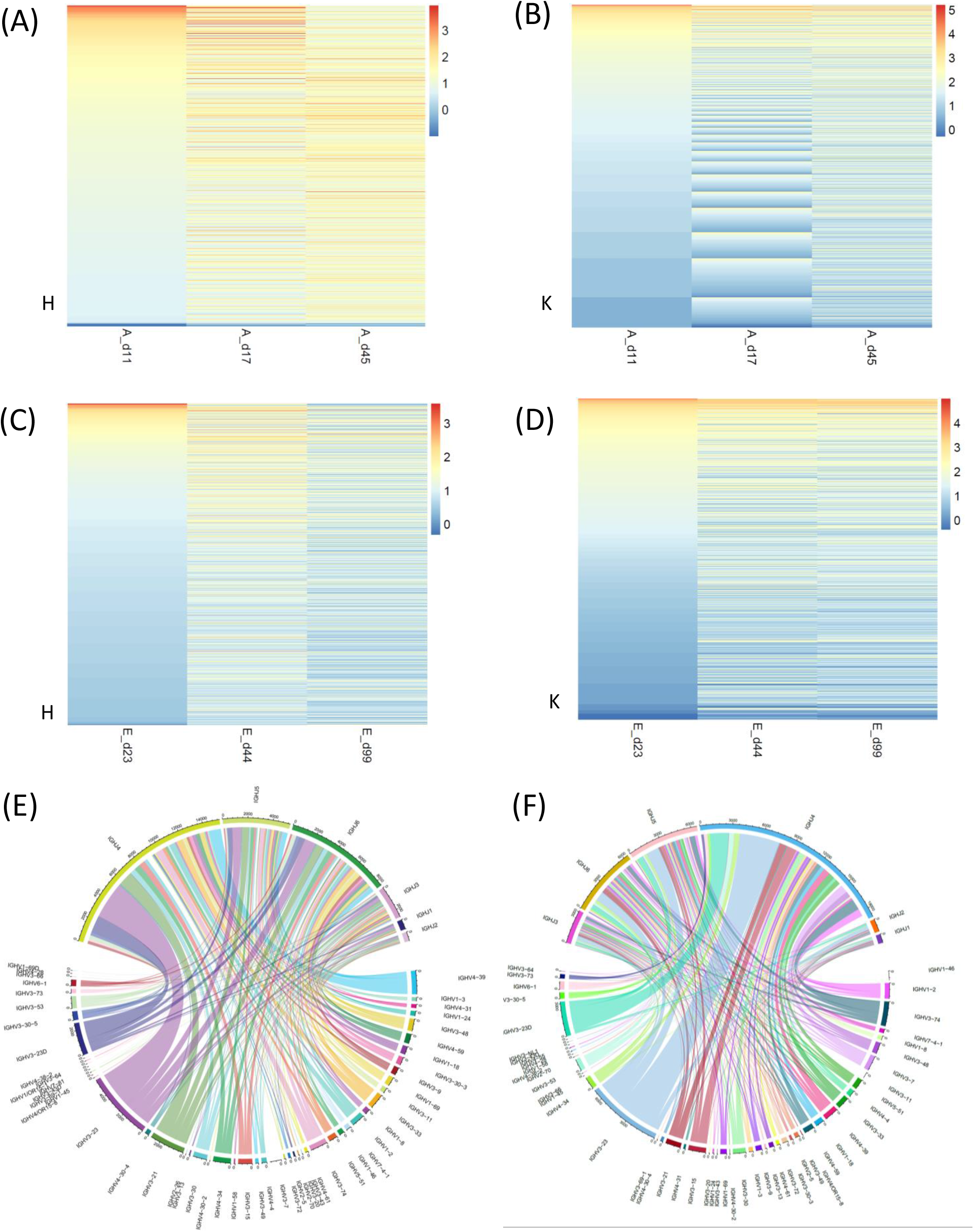
CDR3 sequence based analysis of antibody dynamics. Overlapping public clonotypes across samples were identified based on identical CDR3 amino acid sequence. The total number of sequences (normalized log2 read count) corresponding to the clonotype is represented as a heatmap for heavy chain and light chain for patient A (**A, B**) and patient E (**C, D**). Chord diagram representing V-J segment usage at early timepoint (**E**; day 17) and late timepoint (**F**; day 45) for patient A.

**Figure S4.**
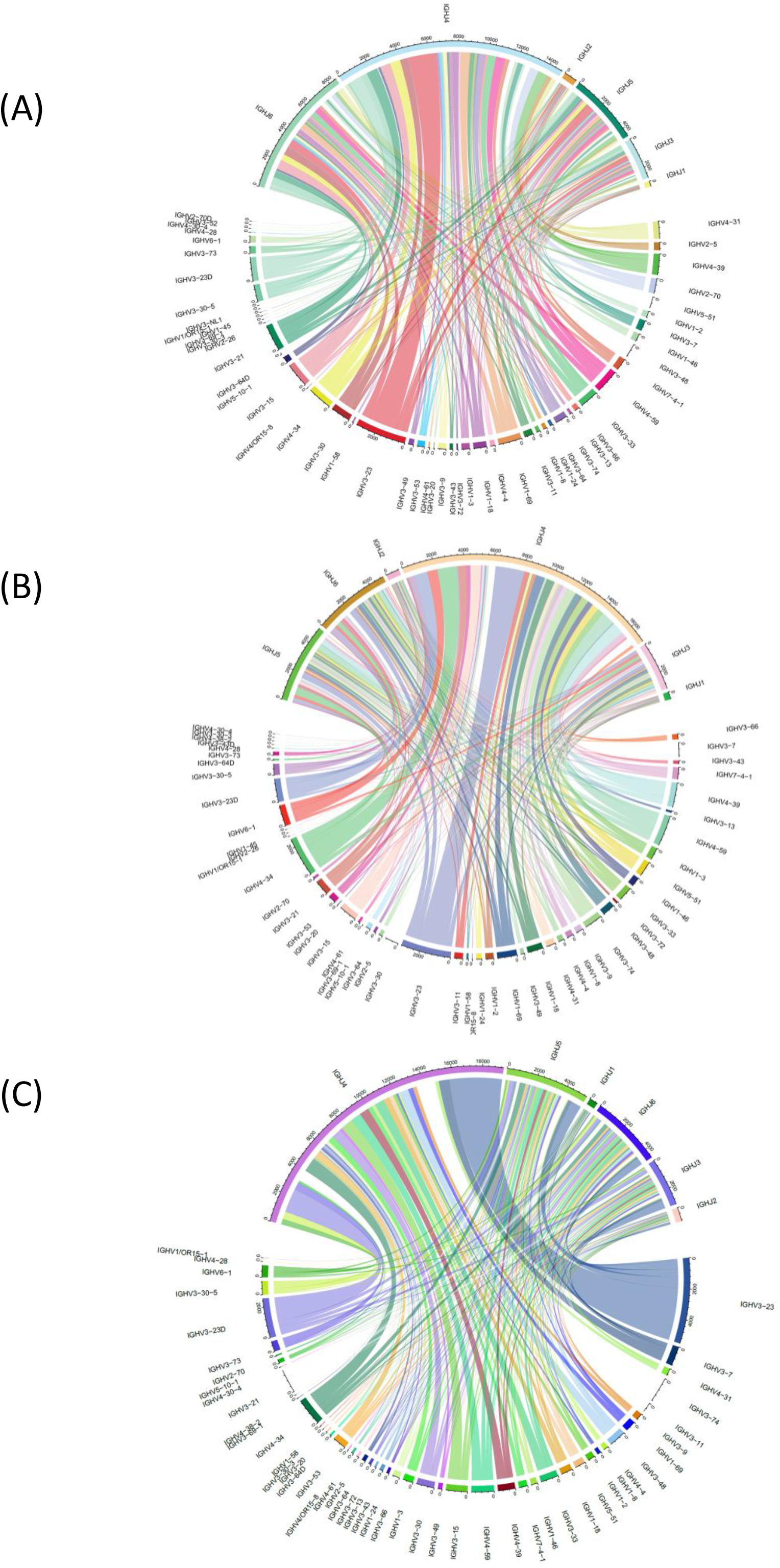
Chord diagram representing V-J segment usage at day 23 (**A**), 44 (**B**) and 99 (**C**) for patient E.

**Figure S5.**
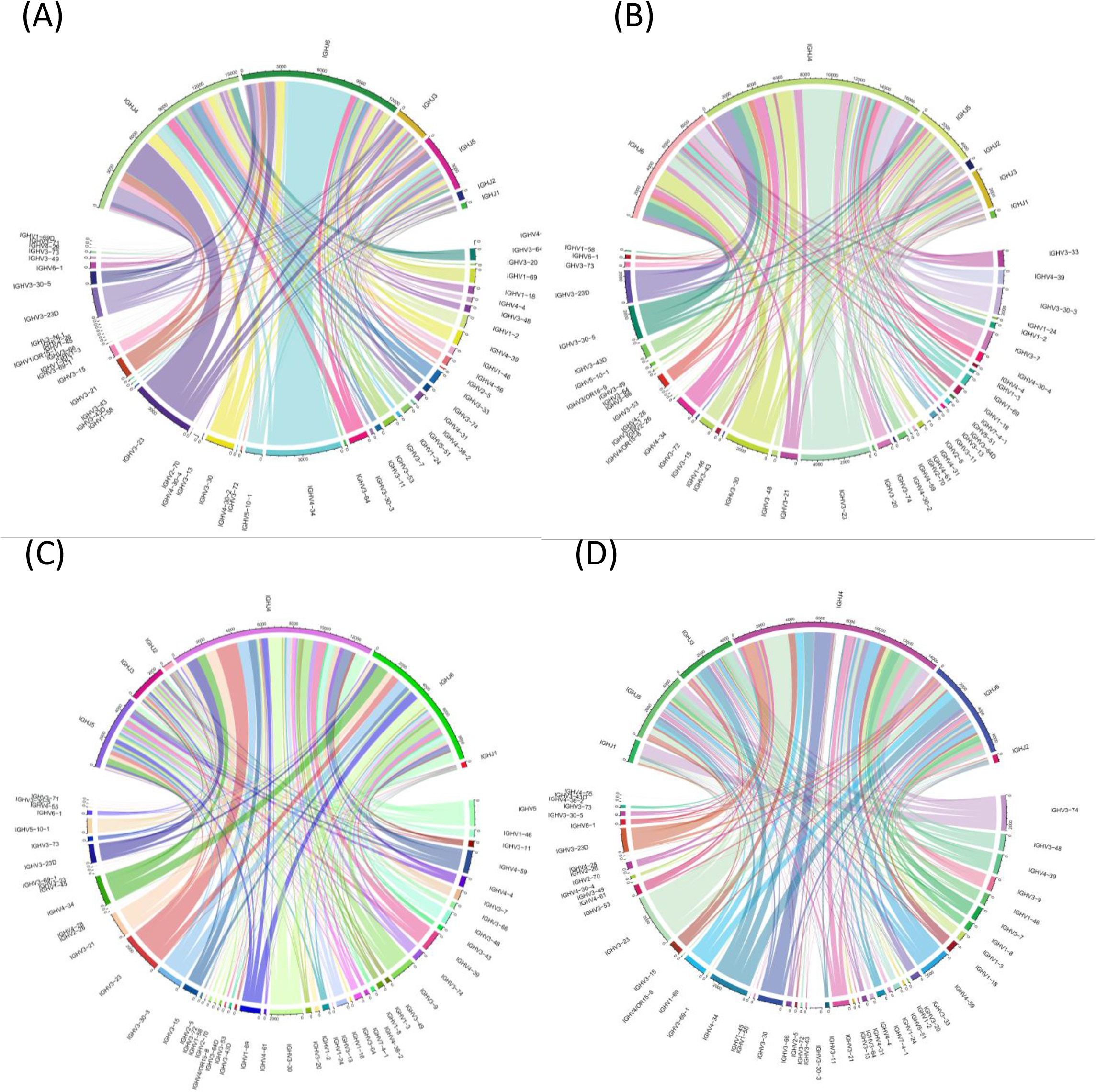
Chord diagram representing V-J segment usage at 2 weeks in mild (Patient C, Patient F) (**A, B**), moderate (Patient P) (**C**) and severe patient (Patient O) (**D**).

